# Locomotory activity is more consistent over time than thigmotaxis and aggressive behavior in sea-ranched Baltic salmon (*Salmo salar* L.)

**DOI:** 10.1101/2025.05.29.656924

**Authors:** Johanna M Axling, Erik Petersson, Svante Winberg

## Abstract

The behavioral differences, individually but also on group level has been evaluated in Baltic salmon parr in a repeated test combining open field and mirror test with focus on aggressive, bold and active behavior. The 164 salmon parr was tested twice and distance moved in arena, activity, was the variable which had the highest correlation between the trials. No correlation was found between aggression and activity or between boldness and activity. The groups which expressed lower aggression had a larger variance in their behavior compared to the highly aggressive groups which had a very strict behavioral repertoire.

## 1 Introduction

Regardless of the animal studied, differences in behavior are present on an individual basis as well as on a group level throughout a range of taxa. When individuals behave in a consistent manner in different situations and over time, this is referred to as animal personality, animal temperament or behavioral syndrome (1–7). The term stress coping style is used when physiological traits are included and the divergent stress coping styles are referred to as proactive and reactive (8,9). Animal personalities have been studied in a large range of species representing mammals, birds, lizards, amphibians, fish, and arthropods (2) in order to evaluate and compare findings across taxa. The so-called bold-shy continuum appears to be present in many species across many taxa (1,10). On one side of this scale there are bold or proactive individuals that tend to be more active and to explore new environments in a superficial but quick manner. Bold individuals react to challenges with active avoidance, aggression and a predominantly sympathetic nervous system activation (9,11,12). At the other side of the continuum there are reactive or shy animals which are usually less active and neophobic but more flexible in their behavioral repertoire. They react to challenges with avoidance and a predominant hypothalamic-pituitary-adrenocortical (HPA) axis activation (the teleostean homolog being the hypothalamic-pituitary-interrenal (HPI) axis) (13). Proactive and reactive individuals have been reported to differ in learning behavior and results from studies on rainbow trout (*Oncorhynchus mykiss*), suggests that reactive individuals are more guided by environmental stimuli whereas the proactive ones more easily form behavioral routines and are less flexible (8,14,15). Similar differences have been observed between bold and shy three spined sticklebacks (*Gasterosteus aculeatus*) regarding foraging behavior, where bolder sticklebacks had a stricter behavioral routine regardless of presence of others compared to shy individuals (16). The authors hypothesized that the cognitive processing which gathers information, recalls past memory, and processes environmental cues could be dependent on how active the individuals are, since that relates to how much area they have had time to cover (16). The authors suggested that the bold and active individuals would process information faster compared to shy individuals which are more reactive to change and their environment and take longer time to process information in a changed situation.

Behavior is a very complex trait since it is not always stable, but instead time and context dependent, and often analyzed by different interconnecting fields, either as a direct target or as a side variable. An additional level of individual variation within the animal personality field is behavioral flexibility or plasticity. Previous classifications of plasticity and flexibility were described by Stamps (17) and categorized in order to develop a framework and to reach consensus in the field. Some of the different forms of plasticity were contextual plasticity (immediate response to current external stimuli), developmental plasticity (learning by life experience), intra-individual variability and endogenous plasticity (behavioral differences within individuals in response to external factors). In order to confirm that a response to a stimulus is a part of an animal’s personality, the research community has relied on demonstration of consistency and repeatability of the response. Meaning that the animal will have to react in a similar manner each time it is exposed to a certain stimulus which may be difficult if the individual has a high level of behavioral plasticity. Coppens et al. (2010) considered behavioral flexibility to reflect the degree to which a behavior is guided by stimuli from the environment, which can be viewed as a vital and potential stable characteristic of coping styles. It is still a matter of debate whether different behavioral profiles or proactive and reactive coping styles show different degrees of behavioral plasticity. Jolles *et al*. (2019) measured on repeated occasions the duration that wild-caught three-spined sticklebacks spent away from a shelter. Shy fish were found to be more plastic in their behavior for the variable exploration without cover and showed the greatest change in the behavior compared to bold individuals. These results seem to confirm the hypothesis that individual variation in boldness is inversely related to behavioral plasticity (18). However, Kim *et al*. (2016)(19) found no link between boldness and plasticity when testing three-spined sticklebacks but a positive relationship between boldness (for fear of novelty or activity after exposure to predatory threat) and plasticity was found by Thomson *et al.* (2012) (20) when testing rainbow trout (*Oncorhynchus mykiss*). Hence it is an ongoing challenge for animal behavior researchers, to form a hypothesis which successfully predicts the relationship between personality and plasticity (21) that holds true for several species and taxa. In addition, some behaviors, such as foraging behavior or exploratory behavior, seem to be easier to repeat compared to other behaviors, such as aggression (22). The reason for this might be that some behavioral traits are more sensitive or dependent on the environment, which might not be stable, or that the behavioral variable is species specific (22).

The aim of the present study was to examine the behavioral consistency over testing occasions of Baltic salmon parr (*Salmo salar L*). In a previous study by Axling *et al.* (23), we found that aggression was not correlated to activity in salmon parr, rather the individuals showing medium to low aggression had high activity. In the current study, the animals were tested twice in a combined open field (OF) and mirror image simulation (MIS) test. We analyzed the consistency over time of three behavioral variables: duration moving in the whole arena (activity), duration in the center zone (boldness) and distance between nose and mirror (aggression). In addition, we tested whether the consistency or plasticity in these parameters was dependent on whether the fish was classified as showing high, medium, low or zero aggression in the first test (23). We hypothesized that highly aggressive individuals would show greater consistency over testing occasions while less aggressive fish would be more plastic in their behavior in subsequent test trials (24–27).

## 2 Material and methods

### 2.1 Animal breeding and maintenance

Experimental protocols and animal handling methods used in this study were approved by the Uppsala Regional Animal Ethics Board (Dnr permit C55/13), following the guidelines of the Swedish Legislation on Animal Experimentation (e.g. Animal Welfare Act SFS 2018:1192) and the European Union Directive on the Protection of Animals Used for Scientific Purposes (Directive 2010/63/EU). The group of fish used in this study presented a subset of an earlier study testing ∼2000 individuals for behavior in the OF and MIS tests (23). In the current study 164 individuals were included, However, not individual could be used due to missing data for some variables.

### 2.2 Behavioral testing

Behavioral trials were conducted from the 30^th^ of August until the 28^th^ October 2016. The first round of trials was conducted from 30^th^ of August to 21^th^ of August and the second round of trials was carried out from the 24^st^ of October to the 28^th^ of October 2016. Fish did not receive feed 24 hours before the trials. The testing arena was a modified white opaque plastic box (Trofast, IKEA, Sweden; dimensions 30 cm length x 42 cm width x 23 cm height, water depth 10 cm) with a mirror on one of the short sides (23). Two detachable hatches were placed parallel to the mirror (23), allowing the experimenter to record both the open field (OF) and mirror stimulation (MIS) tests within the same arena (adapted from 28). The first hatch (29 cm from the mirror) separated the start box from the rest of the arena and the second hatch (8 cm from the mirror) blocked the mirror for the duration of the OF test (5 minutes) of the trial. Unfiltered water from the river Dalälven was use and the temperature ranged from 16.5 (August) to 5.6°C (end of October).

Before the trial the fish were acquired from their home tank, and their PIT tag logged with an ID plate reader (Biomark, USA). The fish was released into the start box of each arena, in total 16 arenas, which were placed in groups of 4 (4 x 4). The trials were video recorded from 4 surveillance cameras (IB8369, Vivotek, New Taipei City, Taiwan), each camera filming 4 arenas. The cameras were mounted from the ceiling in the test room and controlled from the adjacent room. The recording was started directly after the last fish was released into the start box. Five minutes after the recording started the first hatch was removed from all arenas and the OF test was initiated, for a duration of five minutes (minute −5 to 0). The second hatch was removed 10 minutes after the recording started and initiated the MIS test, after which the fish was left undisturbed for the remaining 20 minutes of the trial (minute 0 to 20). The hatches were removed from the arenas by pulling them upwards (which ensured minimal disturbance), using a long rod with a curved edge that hooked into a hole in the gate. The arenas were cleaned with 70% ethanol and rinsed with water between trials. After the first trial was conducted each fish was netted out of the testing arena and anesthetized with Benzocaine solution (50 mg/L) (29). The adipose fin was collected for future biometrics, weight and fork length was recorded. All tested fish were mixed and divided into three new home tanks. For the second trial a new weight and fork length was recorded. After the second trial all fish were randomly released into three home tanks.

### 2.3 Behavioral analysis

The first 5 minutes of each video (before hatch 1 was opened and the fish was in the start box) were disregarded from the analysis. The recorded 25 minutes of each trail (5 min OF and 20 min MIS test) were analyzed using automated tracking software EthoVision XT14 (Noldus Technology, Wageningen, The Netherlands) with nose-tail base detection. Each trial was visually checked for tracking errors and any identified errors were manually corrected. The center zone and the location of the mirror was virtually drawn in the arena (23). We extracted the following three variables from the tracking software for each minute; time spent moving in the whole arena (in cm; activity), time spent in the center zone (in sec; boldness) and distance between nose and mirror when the fish was in the mirror zone (in cm; distance to mirror).

### 2.4 Statistical analysis

The statistical analyses were performed in SAS statistical software version 9.4 (SAS Institute INC, North Carolina, USA). Based on the results of the MIS test from the first trial, each fish was categorized into one of four distinct aggression groups following the procedure described in our previous paper (23). In brief, categorization was based on two different automated tracking variables, the duration of time spent in the mirror zone and the distance moved in the mirror zone which were related to the manually scored variable “striking” (“striking” being defined as a fast movement, resulting from a strong tail beat, towards the mirror at an angle perpendicular to the mirror) using a classificatory discriminant analysis. Aggression groups were highly aggressive (HA), “striking” 30–60 sec/min; low aggressive (LA) 0.001–30 sec/min; zero aggressive (ZA) 0 sec/min. This classification procedure resulted in the following sample size: HA n=22, LA n=68 and ZA n=53. These aggression groups are not related to the variable used for aggression “distance between nose and mirror when fish was in mirror zone”. This differs from the classification used in Axling et al. (23), where four aggression groups were defined (the HA group in the study were split into two groups). However, due to low sample sizes in the HA group, it was not further divided.

Spearman rank correlation coefficients between first and second trial for the three behavioral variables were carried out. The “Total” variable is based on the mean value for all recorded minutes for each individual and “min= −1” is −1 minute before the second hatch was opened. “Min 5” and “Min 10” are 5 and 10 minutes into the MIS test when the mirror is exposed. Water temperature at first and second trial and days between trials were used as covariates.

To obtain the average proportion of time that each individual had, a consistently high value for a behavioral variable for a given trial, the value of a behavioral variable in one minute was correlated to the value in the following minute (Fig S1). For the variables activity (time spent moving in the arena) and boldness (time spent in the center zone) the estimations were done in the following way. Individuals having a value of 40 seconds per minute or higher during both the focal minute and the following minute were classified as having a high level for that variable and minute (Fig S1). The individuals with values between 20 and 40 seconds per minute were classified as medium level and fish having a value of 20 seconds per minute or lower in both minutes were classified as low level. The remaining fish were classified as ‘changers’. For the variable aggression (distance between nose and mirror when the fish was in the mirror zone), the maximum distance was 33 mm. Individuals having a value of 11 mm or less during both the focal minute and the following minute were classified as having a high level for the variable distance to mirror. This was done for all recorded minutes, resulting in 24 comparisons for activity and boldness quantified both during the OF and MIS and 19 values for aggression which was quantified only during the MIS test. From this we calculated the proportion of time (within a trial) that an individual had a high level of a specific behavioral variable; the proportion of time having high activity, proportion of time having high boldness and proportion of time having high aggression. For example, if an individual showed a high level of activity (time spent moving in the arena) for 16 out of the 24 comparisons mentioned above, the proportion having a high activity was calculated as 16/24=0.75. Similar calculations were made for boldness. The results from the second trial were subtracted from the results from the first trial. Negative values mean that the results were greater during the first trial and positive values means that the variable was greater during the second trial.

In the second part of our statistical analysis, we tested whether the above calculated proportions were dependent on whether the fish was classified as showing high, medium, low or zero aggression in the first test (23). We fitted three binomial generalized linear models (i.e. logistic regression) per trial, one for each proportion (proportion displaying high activity, high boldness, or high mirror attraction) using the procedure PROC GENMOD in SAS statistical software. Aggression group was the predictor variable and water temperature, and body weight were added as covariates (the relations between temperature and proportion (probability) of high score is illustrated in Fig. 1A-F). The results from the second trial were subtracted from the results from the first trial. Negative values means that the results were greater during the first trial and positive values means that the variable was greater during the second trial. We evaluated these differences in behavioral consistency for the three different behavioral variables in the different aggression groups, using PROC GLM in SAS statistical software. Levene’s test for homoscedasticity was used due to unequal group sizes and when the assumption of equal variances in aggression groups was violated. For the proportion of time having high boldness, Welch’s ANOVA was used. In all cases Dunn’s post-hoc test was used for pairwise comparisons.

**Fig. 1A.**
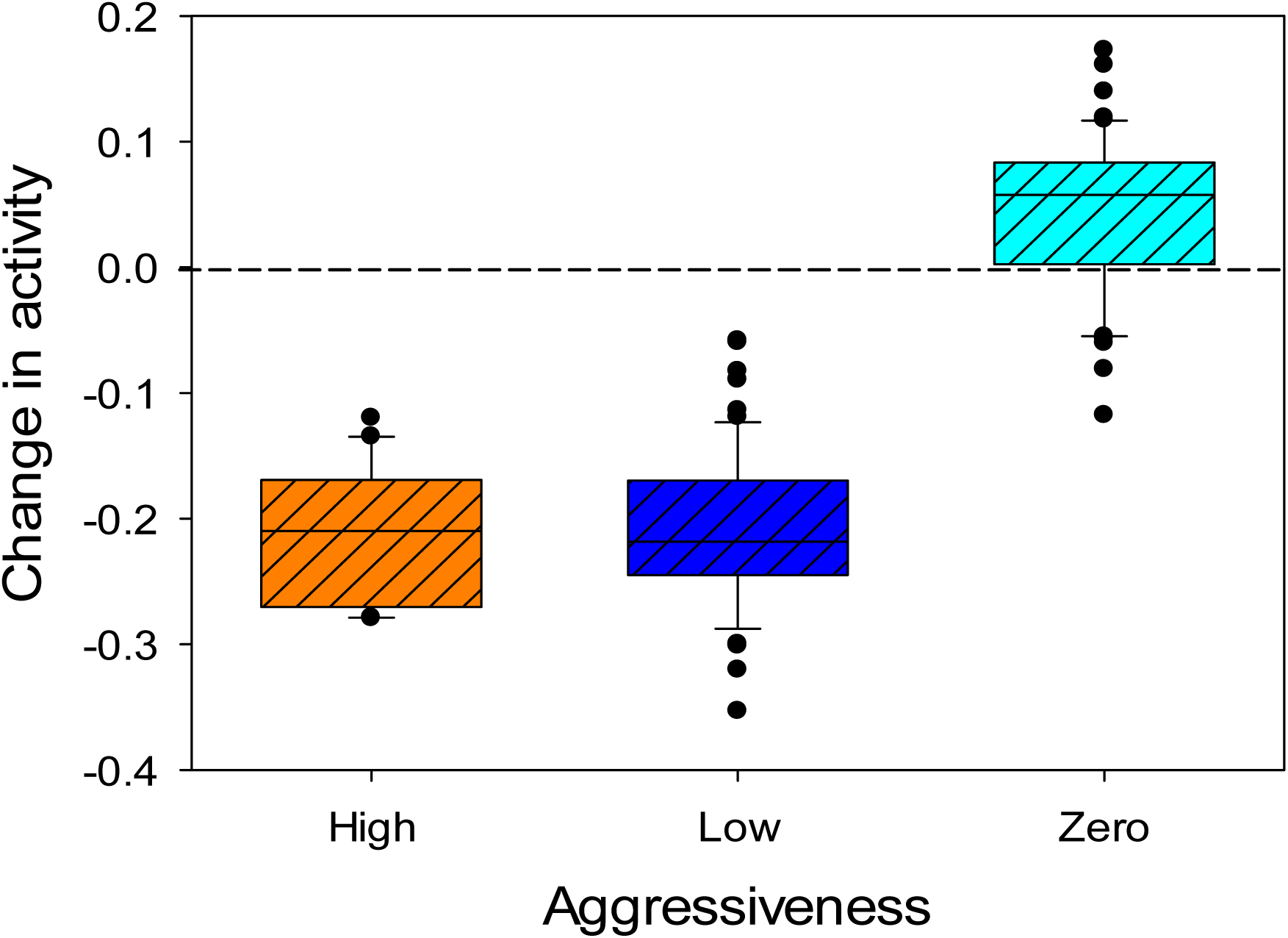
Change in activity (probability of high activity). Levene’s test for homogeneity of variance: F=0.42, p=0.656. ANOVA revealed that the groups differed (F_2,136_=280,0; p<0.001); HA and LA did not differ, and both these two groups differed from ZA at the 5%-level (Dunn’s t-test). In the figure the middle horizontal line in the boxes show the median values, the lower and upper edge of the boxes show the first and third quartile, respectively. The T-bars show the 10^th^ and 90^th^ percentiles, the dots the extreme values.

**Fig. 1B.**
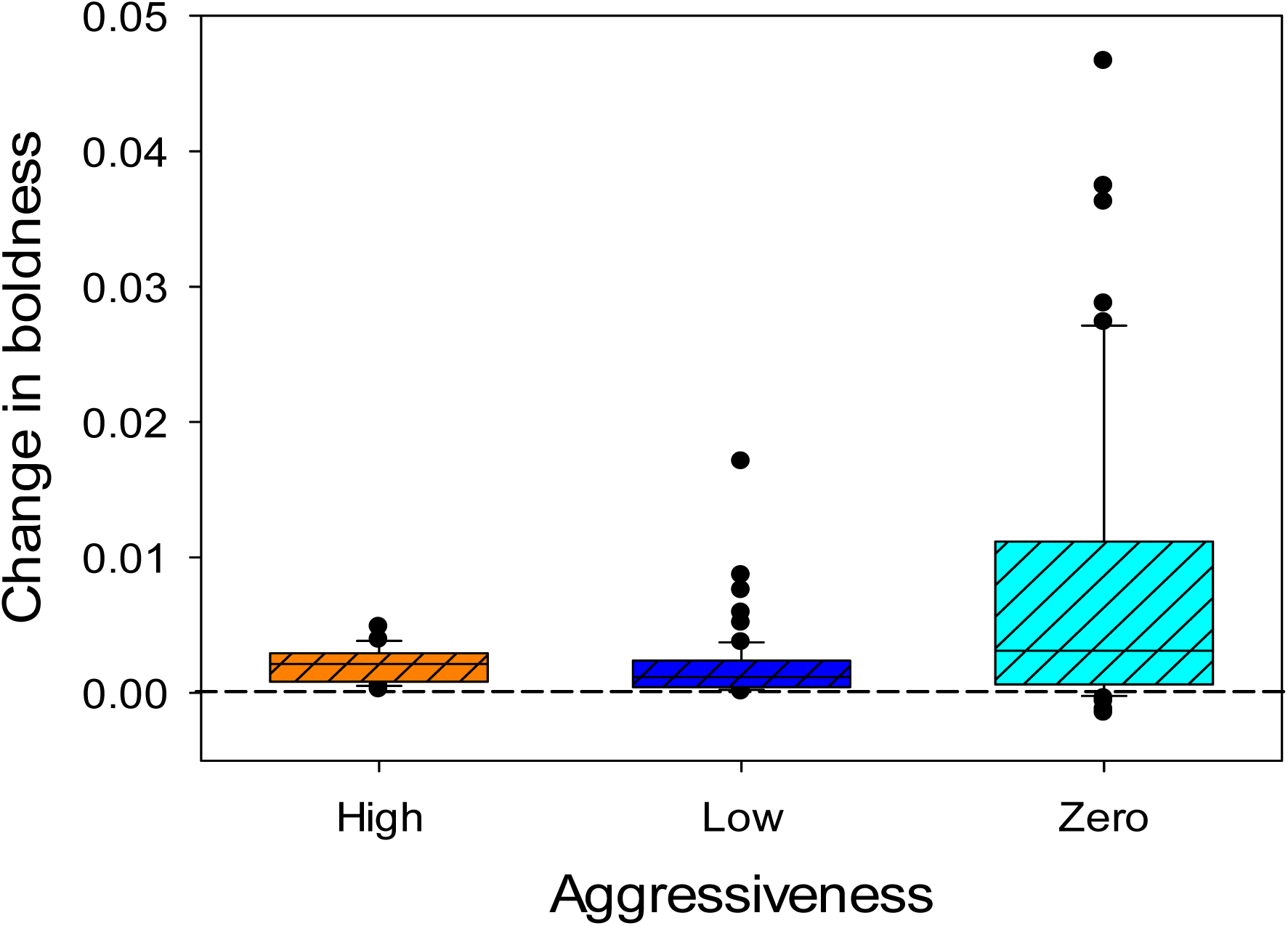
Change in boldness (probability of being in center) Levene’s test for homogeneity of variance: F_2,134_=8.90, p<0.001. According to Welch ANOVA the groups differed (F_2,77.57_=7.63; p<0.001). ZA differed from LA and MA, otherwise there were no significant differences in pair-wise comparisons at the 5%-level (Dunn’s t-test). In the figure the middle horizontal line in the boxes show the median values, the lower and upper edge of the boxes show the first and third quartile, respectively. The T-bars show the 10^th^ and 90^th^ percentiles, the dots the extreme values.

**Fig. 1C.**
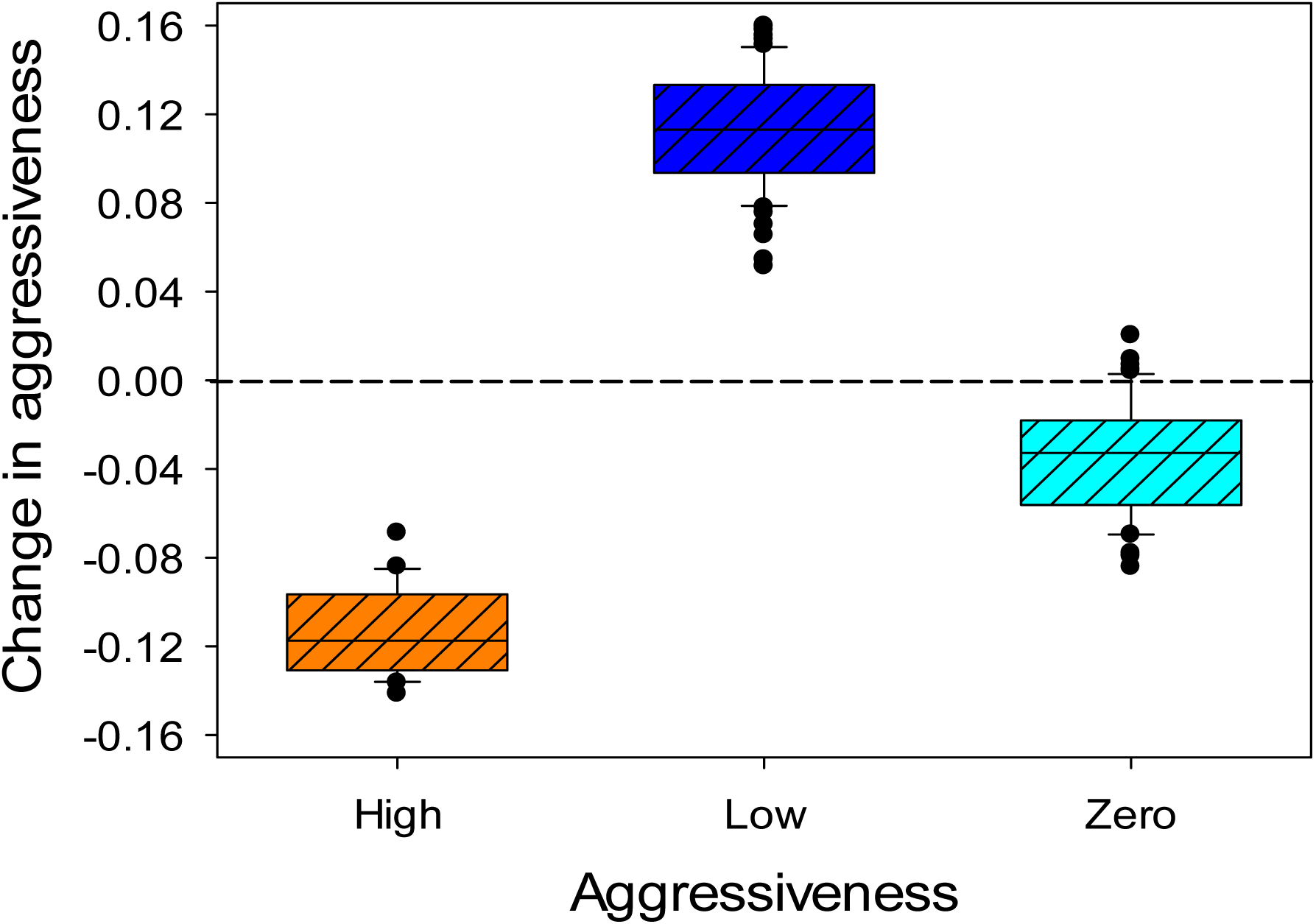
**Change in aggressiveness (probability being close to mirror)**, Levene’s test for homogeneity of variance: F_2,134_=1.52; p=0.222. ANOVA revealed that the groups differed (F_2,136_= 844.9; p<0.001). According to Dunn’s post-hoc test all groups differed at the 5%-level. In the figure the middle horizontal line in the boxes show the median values, the lower and upper edge of the boxes show the first and third quartile, respectively. The T-bars show the 10^th^ and 90^th^ percentiles, the dots the extreme values.

## 3 Results

### 3.1 Individual consistency between the first and second trial

Activity (duration spent moving) showed a positive correlation between tests, with significant correlations between trial 1 and trial 2 for the whole trial, during minute −1, and during minute 10. A trend towards a similar correlation was observed for duration moving during minute 5 (Table 1).

**Table 1:** Spearman rank correlation coefficients between first and second trial for the three behavioral variables. ‘Total’ is based on the mean value for all recorded minutes for each individual. Water temperature at first and second trial and days between trials were used as covariates.

| Variable | Time period (N) | $r_s$ | Raw p-value | Adjusted p-value |
| --- | --- | --- | --- | --- |
| **Activity**<br>Time spent moving in arena (s) | Total (153) | 0.356 | <0.001 | **<0.001** |
|  | min=-1 (153) | 0.284 | <0.001 | **0.002** |
|  | min=5 (152) | 0.196 | 0.017 | 0.065 |
|  | min=10 (153) | 0.305 | <0.001 | **<0.001** |
| **Boldness**<br>Time spent in center zone (s) | Total (153) | 0.188 | 0.021 | 0.081 |
|  | min=-1 (153) | 0.170 | 0.037 | 0.140 |
|  | min=5 (152) | 0.131 | 0.110 | 0.373 |
|  | min=10 (153) | 0.157 | 0.056 | 0.204 |
| **Aggression**<br>Distance between nose and mirror (cm) | Total (153) | -0.005 | 0.951 | 0.999 |
|  | min=-1 (153) | 0.070 | 0.397 | 0.868 |
|  | min=5 (152) | 0.186 | 0.023 | 0.090 |
|  | min=10 (153) | 0.030 | 0.715 | 0.993 |

Boldness (duration in the center zone) did not show a significant correlation between the first and second trial (Table 1). In addition, aggression (distance between nose and mirror) did not show a significant correlation between the first and second trial (Table 1).

### 3.2 Differences between aggression groups

We categorized each individual as displaying high aggression (HA), low aggression (LA) or zero aggression (ZA) on the basis of behavior in the first trial (see Methods section 2.4). From the original variables for activity, boldness and aggression we calculated the proportion of time that each fish displayed a consistent high level of activity, boldness, or aggression over subsequent minutes of each trial (Methods section 2.4). Table 2 shows the results of the logistic regression testing for an effect of aggression group on the proportion of time displaying high activity, boldness, and aggression, whilst controlling for differences in water temperature and body weight. There was a significant difference in the proportion of time having high activity between the aggression groups both for trial 1 and trial 2 (Table 2). During the first trial, there were significant differences between the aggression groups in the proportion of time having high boldness, but the effect of aggression group was no longer significant during the second trial (Table 2). There was a significant difference in the proportion of time having high aggression between the aggression groups for trial 1, but not for trial 2 (Table 2).

**Table 2.**
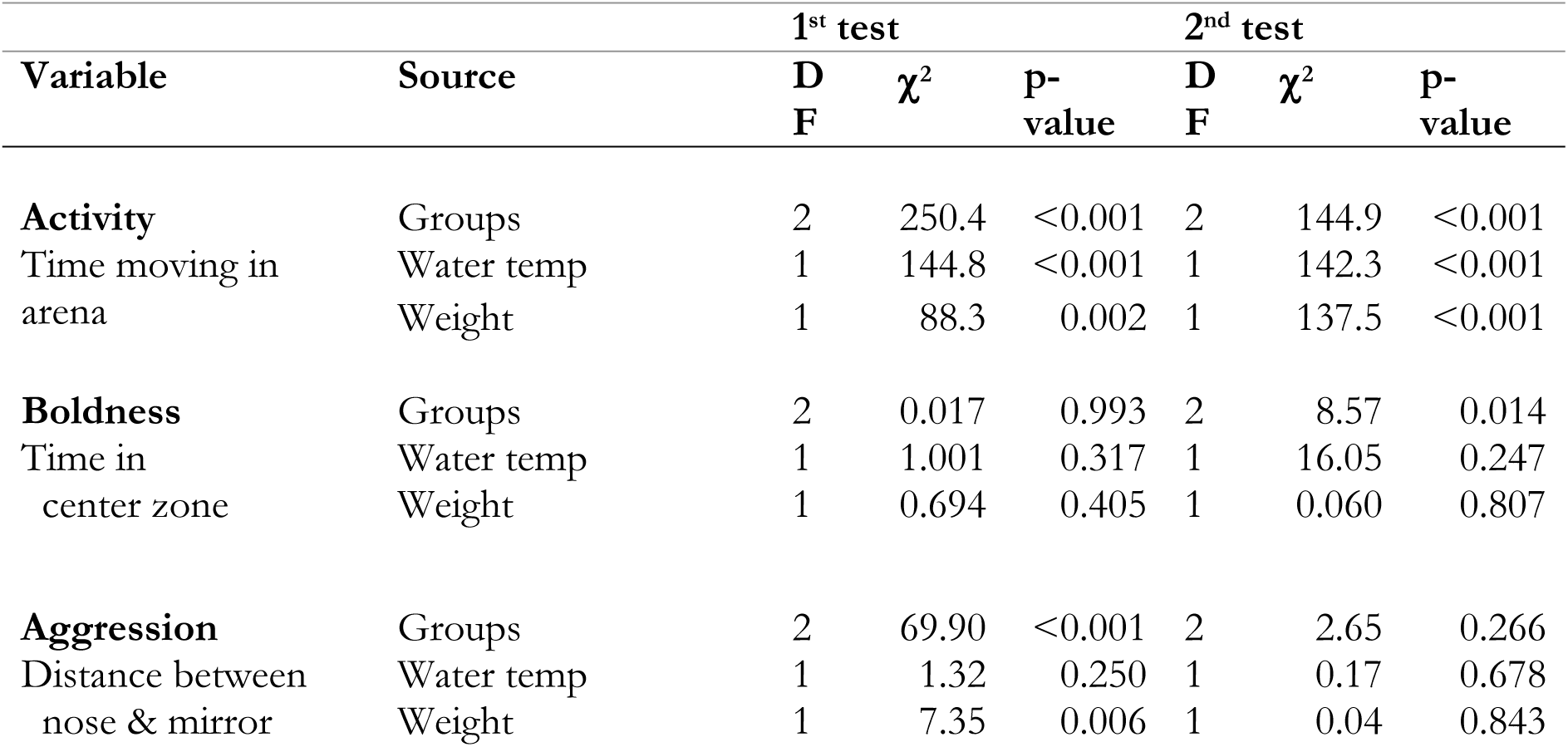
Logistic regression analysis – probability of having high behavioral scores. Total sample sizes; First trial: activity=144; Boldness=144; Aggression=143; Second trial; activity=140; Boldness=137; Aggression=137.

| Variable | Source | 1<sup>st</sup> test |  |  | 2<sup>nd</sup> test |  |  |
| --- | --- | --- | --- | --- | --- | --- | --- |
| | | D<br>F | $\chi^2$ | p-value | D<br>F | $\chi^2$ | p-value |
| **Activity**<br>Time moving in arena | Groups | 2 | 250.4 | <0.001 | 2 | 144.9 | <0.001 |
|  | Water temp | 1 | 144.8 | <0.001 | 1 | 142.3 | <0.001 |
|  | Weight | 1 | 88.3 | 0.002 | 1 | 137.5 | <0.001 |
| **Boldness**<br>Time in center zone | Groups | 2 | 0.017 | 0.993 | 2 | 8.57 | 0.014 |
|  | Water temp | 1 | 1.001 | 0.317 | 1 | 16.05 | 0.247 |
|  | Weight | 1 | 0.694 | 0.405 | 1 | 0.060 | 0.807 |
| **Aggression**<br>Distance between nose & mirror | Groups | 2 | 69.90 | <0.001 | 2 | 2.65 | 0.266 |
|  | Water temp | 1 | 1.32 | 0.250 | 1 | 0.17 | 0.678 |
|  | Weight | 1 | 7.35 | 0.006 | 1 | 0.04 | 0.843 |

During trial 1 the HA and LA group showed the highest activity, and differed significantly from the ZA group, and the same relations were found for trial 2 (Table 3). There was no significant difference in boldness between the aggression groups in trial 1. In trial 2 the ZA group had the highest mean value (Table 3). For aggression, HA and ZA groups had a significantly higher mean than the LA group in the first trial, while in the second trial there was no differences between the aggression groups.

**Table 3.** Posthoc pairwise comparisons between aggression groups and variables for trial 1 and trial 2. Sample sizes within parentheses after the group abbreviation. Mean values denoted with the same letter (column “Difference”) are not significantly different at the 5% level.

| Variable |  | 1st test |  | 2nd test |  |
| --- | --- | --- | --- | --- | --- |
|  | Group | Mean±S.E. | Difference | Mean±S.E. | Difference |
| **Activity**<br>Time spent moving | **HA (20)** | 0.681±0.022 | b | 0.475±0.023 | b |
|  | **LA (65)** | 0.660±0.013 | b | 0.440±0.013 | b |
|  | **ZA (52)** | 0.356±0.014 | a | 0.391±0.014 | a |
| **Boldness**<br>Time in center zone | **HA (20)** | 1.58×10<sup>-14</sup>±4.88×10<sup>-9</sup> | a | 0.0015±0.0015 | a |
|  | **LA (65)** | 1.72×10<sup>-14</sup>±2.92×10<sup>-9</sup> | a | 0.0011±0.0007 | a |
|  | **ZA (52)** | 0.0030±0.0017 | a | 0.0064±0.0025 | b |
| **Aggression**<br>Dist btw nose/mirror | **HA (20)** | 0.678±0.024 | c | 0.563±0.025 | a |
|  | **LA (65)** | 0.445±0.014 | a | 0.559±0.014 | a |
|  | **ZA (52)** | 0.559±0.016 | b | 0.526±0.016 | a |

### 3.3 Differences in behavioral consistency between aggression groups

In order to evaluate the consistency in behavior between the groups, we calculated the difference between trial 1 and 2 in the proportion of time having a high level of activity, boldness and aggression (see Methods section 2.4, Figure 1A-C). The HA and LA group showed the greatest difference in activity from trial 1 to 2, being very active during the first trial but less during the second (Figure 1A). The mean value for all three aggression groups differed from zero (HA group: S=104, p<0.001; LA group: S=-1072.5, p<0.001; ZA: S=458, p<0.001; Signed ranks test, Figure 1A).

All the aggression groups were bolder in the second trial compared to the first (Figure 1B). There were also significant differences between aggression groups in the change in boldness from the first to the second trial. The mean value for all three aggression groups differed from zero (HA group: S=105, p<0.001; LA group: S=1071, p<0.001; ZA: S=622, p<0.001; Signed ranks test, Figure 1B).

The change in aggression from trial 1 to 2 did not differ significantly between aggression groups (Figure 1C). HA and ZA groups were more active during the first trial while the LA group was more active during the second trial. The mean value differed from zero for all three groups (HA: S=-105, p<0.001; LA: S=1072, p<0.001 and ZA: S=-653, p<0.001; Signed rank test, Figure 1C).

### 3.4 Effect of water temperature and body weight

There was a significant effect of water temperature during testing on the proportion of time highly active, both in trial 1 and in trial 2 (Table 2). The activity of all aggression groups increased for lower water temperatures (Figure 2A–B). Proportion of time having high activity was significantly affected by weight for both trial 1 and trial 2 (Table 2). There was no significant effect from water temperature on proportion of time having high boldness (Table 2, Figure 2C– D). During the first trial all aggression groups showed a low probability of being in the center zone, except ZA fish who increased their probability of being in this zone when temperature was above 14°C (Figure 2C). There was no effect from weight on proportion of time having high boldness (Table 2). There was a significant effect from the water temperature on aggression for trial 1 (Table 2, Figure 2E) but not so for trial 2 (Table 2, Figure 2F). Proportion of time having high aggression was significantly affected by weight for trial 1, but not for trial 2 (Table 2).

**Fig. 2.**
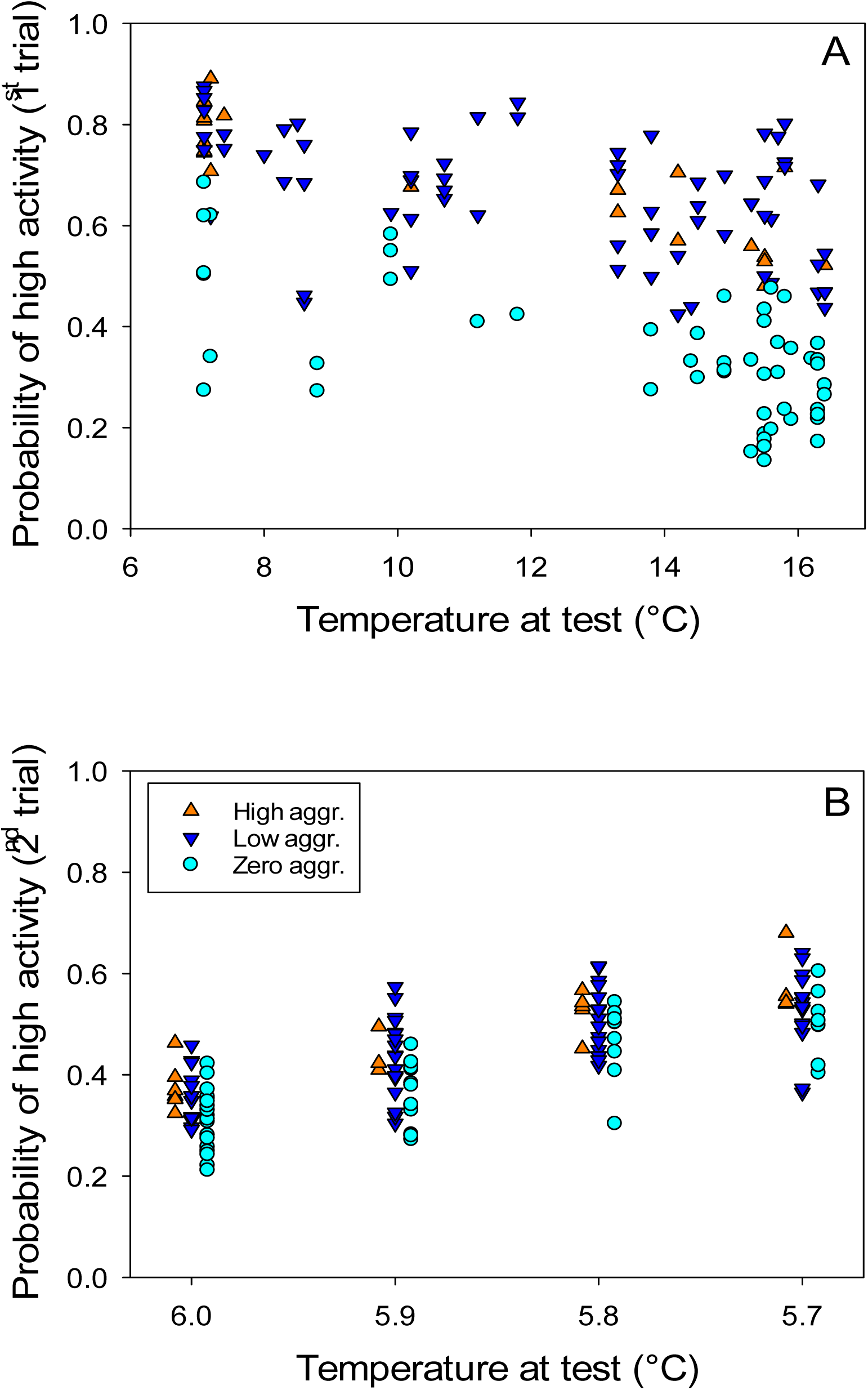
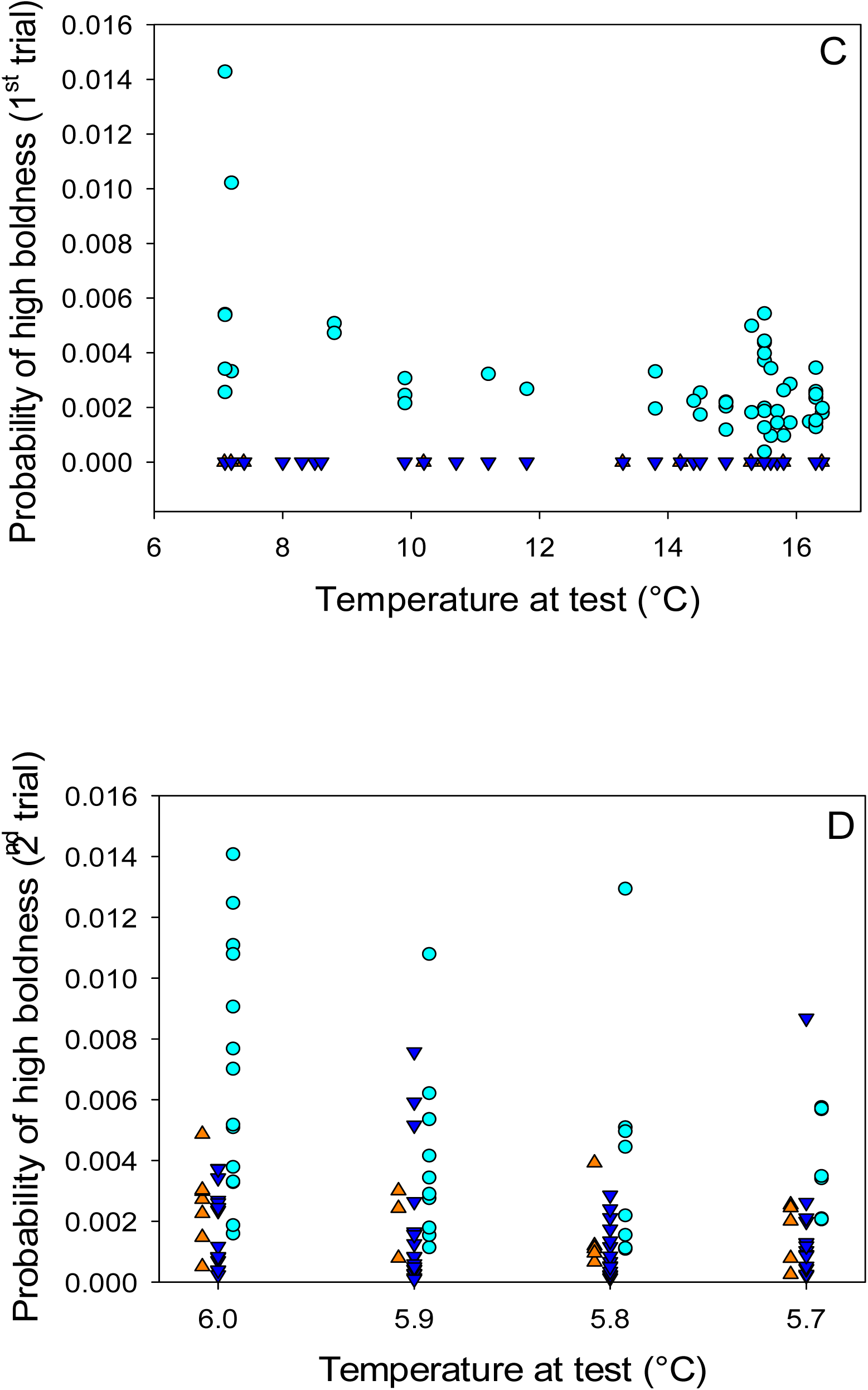
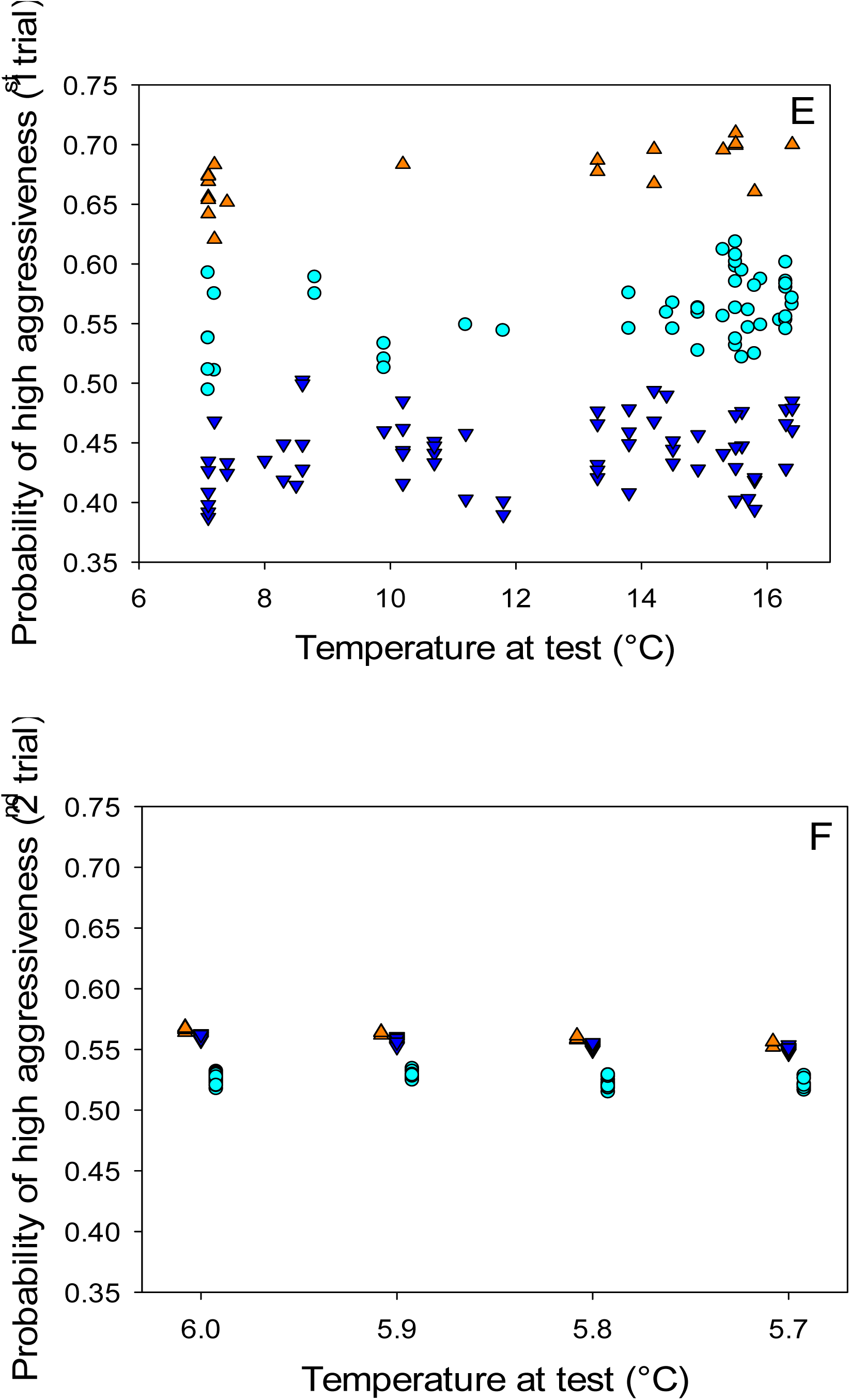
The proportion of time having high activity (A, B) high boldness (C, D) and high aggression (E, F). A, C and E refer to the first trial and B, D and F to the second trial. Please note that the legends are shown only in Fig. 1B – the same legends have been used for all six figures (1A-1F).

## 4 Discussion

In our previous study (23), which used the same behavioral assay as in this, we showed that juvenile Baltic salmon that differ in aggression level also differ in other aspects of their behavioral profile. The most obvious difference was that high and low aggressive fish showed higher levels of activity than high and zero aggression groups. Boldness, as determined by willingness to visit and spend time in the center of the arena, was correlated with activity. However, aggression was not correlated to either activity or boldness. In the current study, we investigated the consistency over two testing occasions for activity, boldness and aggressive behavior. In addition, we analyzed whether consistency over time was related to aggression level.

The results show that activity is the behavioral variable which is the most stable between trials. The time spent moving showed relatively strong correlation between test one and test two whereas no such correlations were observed for time spent in the center zone (boldness) or distance between nose and mirror (aggression). Similarly, in brown trout (*Salmo trutta*) individual differences in activity in an open field test was the most consistent trait between test sessions (28). However, in their study activity in the novel object test was also consistent between tests whereas activity in mirror tests was not. Moreover, in the study by Adriaenssens and Johnsson (28) activity was found to correlate with both aggression and boldness forming a behavioral syndrome. In our previous study we showed that neither activity nor boldness were correlated to aggression in this population of Baltic salmon, even though activity and boldness were (23). Predation is known to be important for the development of behavioral syndromes (30,31). In fact, Bell and Sih (2007) (32) showed that sticklebacks subjected to predation in a laboratory setting developed a stronger correlation between aggressiveness and boldness. In the study by Adriaenssens and Johnsson (28) the fish were wild caught and thus subjected to predation during their early development. During early development when fry emerges from the gravel, both salmon and brown trout are subjected to a strong selection bottleneck with very high mortality (33,34). Hatchery reared fish, as the salmon used in our studies, do not experience any predation and show much lower early mortality. The lack of predation and divergent selection in the hatchery environment could be factors affecting the development of consistent behavioral profiles.

Even though aggression was not consistent between the tests we found that the consistency of other behavioral variables over trials differed between salmon displaying different aggression levels, where fish displaying a low level of aggression were less consistent in their behavior than highly aggressive fish. This result suggests an inverse relationship between behavioral plasticity and aggressiveness. Overall, aggression group was a strong predictor of behavioral variables, even though water temperature and weight also affected the behavior.

Classification of the fish in aggression groups was based on behavior during the first trial with fish classified as highly aggressive showing highest interest in the mirror. However, the interest in the mirror declined during the second trial. When tested in the second trial, the behavioral arena was not novel to the fish, even though the second trial was performed 1–8 weeks after the first trial. Thus, the process of habituation not only during a trial but also between trials may make repeated testing difficult (35).

The fact that handling and repeated testing is stressful to the fish adds to the difficulties and reliability of repeated testing. In this study all tested fish were mixed after their first trial and transferred to new holding tanks. The handling of the fish in connection with behavioral testing most likely resulted in additional stress reactions. The transfer to new holding tanks, mixing fish from previous groups, will have resulted in the development of new dominance hierarchies, which may be very stressful as well. Stress and social interaction may have large behavioral effects and personality traits appear to be much more plastic than originally realized (36). For instance, when the HR and LR strains of rainbow trout, which were selected for high and low post-stress plasma cortisol respectively, were transported from Windermere to Oslo, the behavioral profiles were totally reversed with the reactive HR trout acclimating faster than the proactive LR and becoming dominant in dyadic fights more often than LR trout (14). Thus, traits like boldness and aggressiveness may not be fixed and individuals may not have a permanent phenotype. Several studies in rainbow trout investigating plasticity in boldness have shown that the dynamic nature of bold and shy traits. When given the experience of winning dyadic fights, shy trout increased their boldness, whereas bold trout losing fights became shyer (37). Moreover, Thomson et al. (20) showed that when bold trout were transferred to groups containing either 100% bold or shy individuals, they became shyer, but shy trout transferred in a similar way remained shy. However, physiological traits such as HPI axis reactivity appear to be more stable and less affected by the social environment. In the study by Thomson et al. (20) trout originally classified as bold continued to show lower post-stress plasma cortisol than shy trout even after transfer to new groups. Similarly, in HR and LR trout still differed in HPI axis reactivity, HR showing higher post-stress plasma cortisol than LR trout, even after transport in spite of the dramatic shift in behavioral profiles (14). Additional studies have found a low behavioral change in extreme behaviors. For instance, an earlier study showed that aggressive mice did not change their level of aggression across social context, whereas less aggressive mice did (38).

When investigating repeatability in behavior it is also important to consider the state and development of the individual since these factors may greatly affect the behavioral reaction and outcome of the test. The fish tested in our study were most likely in a similar state at both trials (developmental stage, size, age, and condition). Salmon has a complicated and plastic life history. The river Dalälven population smoltifies at an age of 1–2 years (39). Moreover, a relatively large proportion of the males get sexually mature as parr, at a similar age (39). Smoltification as well as sexual maturation have large effects on the behavior, precociously mature males being bold and aggressive (40) whereas smoltifying fish show reduced aggression (41). The fish used in the present study were only 7 months old which is considered too early for sexual maturation or smoltification (42). However, some physiological differences related to the divergent developmental trajectories might already have been initiated in some individuals.

In conclusion, our results show that juvenile Baltic salmon classified according to individual levels of aggressive behavior differ also in other behavioral traits. Still, in most cases behavioral profiles appear plastic and differences in behavioral traits were not stable between the two trials, the exception being activity. However, for low aggressive fish activity was more variable between the trials than for high aggressive fish, suggesting that low aggressive fish are more plastic in their behavioral repertoire than high aggressive fish. Body mass as well as water temperature had effects on the behavior of the fish. Thus, it cannot be excluded that the shift in behavioral profiles between the trials is also related to the growth and development of the fish. Even though the fish had not reached the age where smoltification or male sexual maturation is expected, the fish may have entered different developmental trajectories leading to these divergent life histories, i.e. smoltification and anadromy or precautious sexual maturation, respectively.

## Acknowledgment

This study was financed by the Swedish Research Council (to SW) and Facias foundation (to JA). The authors thank Uppsala University Behavioural Facility (UUBF), Disciplinary Domain of Medicine and Pharmacy, Uppsala University. We are also grateful for technical assistance by the staff at Älvkarleby hatchery (Swedish Agricultural University). We would also like to thank Laura Vossen, Airon Liun and Fredrik Axling for great work effort related to the project and Prof. Erika Roman for valuable comments on the manuscript.

## Author contributions

JA tested the fishes, interpreted the statistical analyses, and wrote the first draft of the article. EP conducted the statistical analyses. SW helped in writing the manuscript and supervised the experimental design. All authors contributed to manuscript revision, read, and approved the submitted version.

## For Supplementary Material

**Fig. S1.**
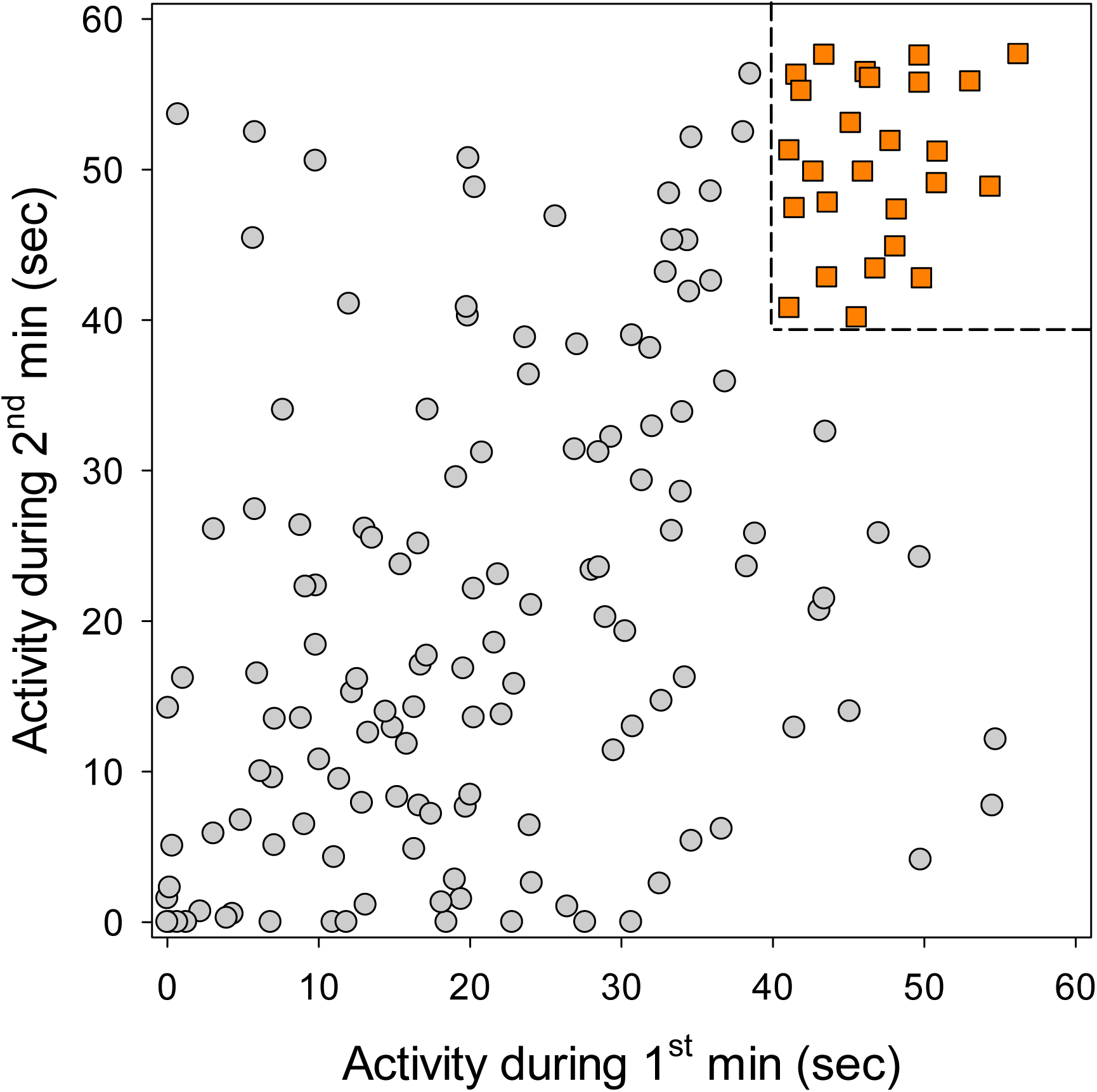
The figure shows the activity level of the investigated individual of juvenile Atlantic salmon (*Salmo salar* L.) during the first minute and the second minute of the experiment. The activity level was retrieved using Ethovison® software and the software recorded the number of seconds (0–60) an individual was moving (swimming). The values at the upper right (orange squares) show the individuals that had high activity during both the first and second minute of the experiment.

Similar comparisons were made for the second vs the third minute, the third vs the fourth minute and so on up the 24^th^ vs the 25^th^ minute, giving 24 figures as the figure above.

For each individual we calculated the number of times each individual had been in the upper right area (i.e. having high activity). As an example, if an individual had been registered in the upper right area 12 out of the 24 possible comparisons, the proportion (probability of) high activity was 12/24=0.5.

## Notes

### Competing Interest Statement

The authors have declared no competing interest.

